# Influence of hyaluronic acid transitions in tumor microenvironment on glioblastoma malignancy and invasive behavior

**DOI:** 10.1101/291898

**Authors:** Jee-Wei E Chen, Sara Pedron, Peter Shyu, Yuhang Hu, Jann N Sarkaria, Brendan A C Harley

**Affiliations:** Department of Chemical & Biomolecular Engineering, University of Illinois at Urbana-Champaign, Urbana, IL, USA; Carl R. Woese Institute for Genomic Biology, University of Illinois at Urbana-Champaign, Urbana, IL, USA; Department of Mechanical Science and Engineering, University of Illinois at Urbana-Champaign, Urbana, USA; Department of Radiation Oncology, Mayo Clinic, Rochester, MN, USA

**Keywords:** cell invasion, hyaluronic acid, hydrogels, tumor microenvironment, tumor margins, molecular weight, glioblastoma, brain tumor

## Abstract

The extracellular matrix (ECM) is critical in tumor growth and invasive potential of cancer cells. In glioblastoma tumors, some components of the native brain ECM such as hyaluronic acid (HA) have been suggested as key regulators of processes associated with poor patient outlook such as invasion and therapeutic resistance. Given the importance of cell-mediated remodeling during invasion, it is likely that the molecular weight of available HA polymer may strongly influence GBM progression. Biomaterial platforms therefore provide a unique opportunity to systematically examine the influence of the molecular weight distribution of HA on GBM cell activity. Here we report the relationship between the molecular weight of matrix-bound HA within a methacrylamide-functionalized gelatin (GelMA) hydrogel, the invasive phenotype of a patient-derived xenograft GBM population that exhibits significant in vivo invasivity, and the local production of soluble HA during GBM cell invasion. Hyaluronic acid of different molecular weights spanning a range associated with cell-mediated remodeling (10, 60, and 500 kDa) was photopolymerized into GelMA hydrogels, with cell activity compared to GelMA only conditions (-HA). Polymerization conditions were tuned to create a homologous series of GelMA hydrogels with conserved poroelastic properties (i.e. shear modulus, Poisson’s ratio, and diffusivity). GBM migration was strongly influenced by HA molecular weight. While markers associated with active remodeling of the HA content, hyaluronan synthase and hyaluronidase, were found to be insensitive to matrix immobilized HA content. These results provide new information regarding the importance of local hyaluronic acid content on the invasive phenotype of GBM.

## 1. Introduction

Glioblastoma (GBM), a WHO grade IV astrocytoma, is the most common and deadly form of brain cancer and accounts for more than 50% of primary brain tumors (Furnari et al., 2007;Nakada et al., 2007;Wen and Kesari, 2008). Unlike many other cancers that metastasize to a secondary site, GBM instead is known to diffusely infiltrate throughout but rarely metastasize beyond the brain, and this invasive phenotype contributes to poor patient prognosis (median survival < 15 months and 5 year survival < 5%) (Stupp et al., 2005;Jackson et al., 2011;Johnson and O’Neill, 2012;Mehta et al., 2015). The brain extracellular matrix and GBM tumor microenvironment (TME) display striking differences to other tumors, show a large amount of spatial and temporal heterogeneity, and can differ patient-to-patient. However, while fibrillar proteins such as collagen and fibronectin are abundant in many other tissues, the brain ECM has minimal fibrillar structures and is mainly composed of hyaluronic acid (HA, also called hyaluronan or hyaluronate) (Bonneh-Barkay and Wiley, 2009;Sivakumar et al., 2017).

The GBM TME is not homogeneous but a complicated heterogeneous environment, especially on the tumor margins, where transitions between the tumor microenvironment and surrounding brain parenchyma are characterized by transitions in structural, biomolecular, and cellular composition. The matrix compositional transition from natural brain to tumor provides a potential invasion path for GBM and, therefore, might contribute to poor patient prognosis (Syková, 2002;Quirico-Santos et al., 2010;Charles et al., 2011;Jackson et al., 2011;Wiranowska and Rojiani, 2011;Junttila and de Sauvage, 2013). Processes of GBM invasion, particularly in the perivascular niche in the tumor margins, involve exposure to not only HA but a range of fibrillar protein content and significant matrix remodeling, resulting in GBM cell exposure to not only HA but also a wide range of molecular weights of HA (Bayin et al., 2014;Lathia et al., 2015;Paw et al., 2015). In this context, the amount and molecular weight distribution of HA, associated with constant turnover from oligosaccharides to high MW HA, across the tumor microenvironment is believed as an important regulator of GBM invasion (Itano and Kimata, 2008). Hyaluronic acid, a negatively charged, nonsulfated GAG, is the main component of brain ECM. HA is naturally produced by hyaluronan synthase (HAS) family and degraded by hyaluronidase (HYAL) in mammalian animals (Misra et al., 2011). While the presence of HA has been shown to be important to tumor progression (Toole, 2004;Stern, 2008;Kim and Kumar, 2014), significant investigation is needed to explore the role of the molecular weight (MW) of HA on processes associated with GBM invasion, progression, and therapeutic response.

Remodeling of hyaluronic acid in the context of GBM cell invasion requires the combined effort of a range of degradative and biosynthetic proteins. Notably, HA biosynthesis is driven by hyaluronic synthase (HAS), which has multiple isoforms responsible for secreting different MW HA (HAS1: 200-2000kDa; HAS2: >2000kDa; HAS3: 100-1000kDa). Similarly, the degradation of HA by hyaluronidase (HYAL) can produce final fragments with different MW. In GBM, HYAL1 (<20kDa) and HYAL2 (20-50kDa) are the most abundant HYAL isoforms (Misra et al., 2011;Khaldoyanidi et al., 2014). Due to the constant synthesis and degradation of HA, a wide range of different molecular weight HA (High, >500kDa; Medium, 50-350 kDa; Low, <30 kDa) are present in the brain and TME (Toole, 2004;Lam et al., 2014;Monslow et al., 2015). HMW HA is important for structural support and the biophysical properties in tissue, and is directly synthesized via HAS. While HMW HA can inhibit tumor growth in colon cancer (Mueller et al., 2010) it also decreases production of MMPs by suppression of MAPK and Akt pathways (Chang et al., 2012). L-MMW, generated from HYAL degradation as final products, are often associated with enhanced invasion and increased tumor growth (Monslow et al., 2015). LMW and MMW HA have been reported to enhance cancer proliferation, cell adhesion as well as secretion of MMPs for matrix remodeling (Tofuku et al., 2006). LMW HA has also been reported to be pro-inflammatory and pro-angiogenic, which may contribute to cancer invasion (West et al., 1985;Lam et al., 2014). In contrast, the effects of oligo HA have been more variable. In papillary thyroid carcinoma, oligo HA is associated with increased (Dang et al., 2013), while other studies demonstrate suppression of signaling pathways such as Ras and Erk and reduced tumor progression (Misra et al., 2006;Toole et al., 2008).

Despite the conflicting HA-cancer relations and lack of full understanding of HA MW contribution, HA clearly plays a significant role in many signaling pathways and in tumor progression. In this study, we analyze the effects of matrix-bound HA on GBM cell invasion by using an in vitro fully three-dimensional gelatin based hydrogel system that our lab has previously developed (Pedron et al., 2013b;Chen et al., 2017c). Previous efforts have used this platform to demonstrate the effect of a single MW HA immobilized within the GelMA hydrogel on the invasive phenotype of GBM cell lines as well as the gene expression signature and response to a model tyrosine kinase inhibitor (erlotinib) (Chen et al., 2017a;Pedron et al., 2017a;Pedron et al., 2017b;Chen et al., 2018). Here we selectively decorate the GelMA hydrogel with a range of MW HA spanning those seen in the GBM TME (10, 60, 500 kDa). Further, we examine the behavior of a patient-derived xenograft (PDX) GBM specimen that maintains patient specific molecular and morphologic characteristics (Sarkaria et al., 2006;Sarkaria et al., 2007). We evaluate cell growth, invasion, and proteomic responses of GBM cells within our platform and demonstrate the influence HA MW on GBM invasive phenotype.

## 2. Materials and Methods

### 2.1. Hydrogel fabrication and characterization

Fabrication of methacrylated gelatin (GelMA) and methacrylated hyaluronic acid (HAMA) precursors and hydrogels were as described in previous publications (Pedron et al., 2013b;Chen et al., 2017c). Briefly, gelatin powder (Type A, 300 bloom from porcine skin, Sigma-Aldrich) was dissolved in 60°C phosphate buffered saline (PBS; Lonza, Basel, Switzerland) then methacrylic anhydride (MA; Sigma-Aldrich) was added into the gelatin-PBS solution dropwise and allowed the reaction proceed for 1 hour. The GelMA solution was then dialyzed (12-14 kDa; Fisher Scientific) and lyophilized. HAMA was synthesized by adding 10 mL MA dropwise into a cold (4°C) HA sodium salt (10, 60 or 500 kDa; Lifecore Biomedical) solution (1g HA sodium salt in 100 mL DI water). The pH was adjusted to 8 with the addition of 5N sodium hydroxide solution (NaOH; Sigma-Aldrich) and the reaction proceeded overnight at 4°C. The product was then purified by dialysis and lyophilized. The degree of MA functionalization of both GelMA and HAMA was determined by ^1^H NMR (data not shown) (Pedron et al., 2013b;Chen et al., 2017c).

Hydrogels (GelMA ± HAMA) were prepared by dissolving GelMA and HAMA in PBS at a total concentration of 4 wt% with gentle heating (37°C ∼45°C) in the presence of a lithium acylphosphinate (LAP) as photoinitiator (PI, adjusted to maintain same Young’s modulus). The mixture was placed into Teflon molds (0.15 mm thick, 5 mm radius) and photopolymerized under UV light (AccuCure LED 365 nm, Intensity 7.1 mW/cm^2^) for 30s (Mahadik et al., 2015). Cell-containing hydrogels were made similarly but with addition of cells (4×10^6^ cells/ mL hydrogel solution) to the pre-polymer solution, prior to pipetting into Teflon molds, and then photopolymerized. Details regarding the hydrogel compositions are listed in Table 1. All HA containing GelMA hydrogel groups were fabricated with 15% w/w HA, consistent with previous HA-decorated GelMA hydrogels described by our group (Pedron et al., 2015;Chen et al., 2017a;Pedron et al., 2017a;Pedron et al., 2017b;Chen et al., 2018).

**Table 1.** Hydrogel composition and characterization results (n=6).

| **Hydrogel** | **-HA** | **10K** | **60K** | **500K** |
| --- | --- | --- | --- | --- |
| **GelMA (wt%)** | 4.0 | 3.4 | 3.4 | 3.4 |
| **HAMA (wt%)** | 0 | 0.6 | 0.6 | 0.6 |
| **HA sodium salt MW** | N/A | ~ 10 kDa | ~ 60 kDa | ~ 500 kDa |
| **LAP (wt%)** | 0.1 | 0.02 | 0.02 | 0.02 |
| **Young's Modulus (kPa)** | $2.76 \pm 0.24$ | $2.97 \pm 0.15$ | $2.79 \pm 0.15$ | $2.70 \pm 0.03$ |
| **Diffusivity ($\mu\text{m}^2/\text{s}$)** | $161.04 \pm 70.33$ | $153.54 \pm 34.92$ | $169.90 \pm 26.88$ | $156.43 \pm 50.18$ |

### 2.2. Characterization of hydrogels

#### 2.2.1. Young’s modulus

The compressive modulus of each hydrogel variant was measured using an Instron 5943 mechanical tester. Hydrogels were tested under unconfined compression with a pre-load 0.005N at the rate of 0.1 mm/min, with their Young’s modulus obtained from the linear region of the stress-strain curve (0-10 % strain).

#### 2.2.1. Diffusivity

The water diffusivity of each hydrogel was measured through indentation tests using atomic force microscopy (AFM, MFP-3D AFM, Asylum Research) (Figure 1). The stiffness of the cantilever used in the measurements is 0.6 N/m. A spherical polystyrene probe of 25 μm diameter was attached to the tip (Novascan). Three separate measurements of different indentation depths were taken. After surface detection, the spherical indenter was pressed into the sample to a certain depth in the rate of 50 μm/s and was held for a period of time until the force on the indenter reaches a constant value. The force on the indenter was measured as a function of time F(t). The time-dependent response of hydrogels is due to solvent migration. The poroelastic relaxation indentation problem has been solved theoretically by Hu *et al.* (Hu et al., 2010;Hu et al., 2011). Simple solutions have been derived for direct extraction of material properties from the relaxation indentation measurement. According to this method, the normalized force relaxation function is a function of a single variable: the normalized time *τ* = *Dt*/*a*^2^, with D being the diffusivity, t being time, and a being the contact radius that is related to the radius of the spherical probe R and indentation depth h by 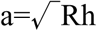:

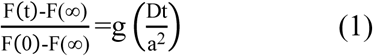

This master curve has been derived numerically as

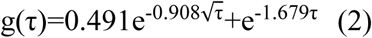

Normalizing the experimental data and fitting it with the theoretical curve (Eq.2), we can extract the single fitting parameter diffusivity D. More details can be seen in references (Hu et al., 2010;Hu et al., 2011).

**Figure 1.**
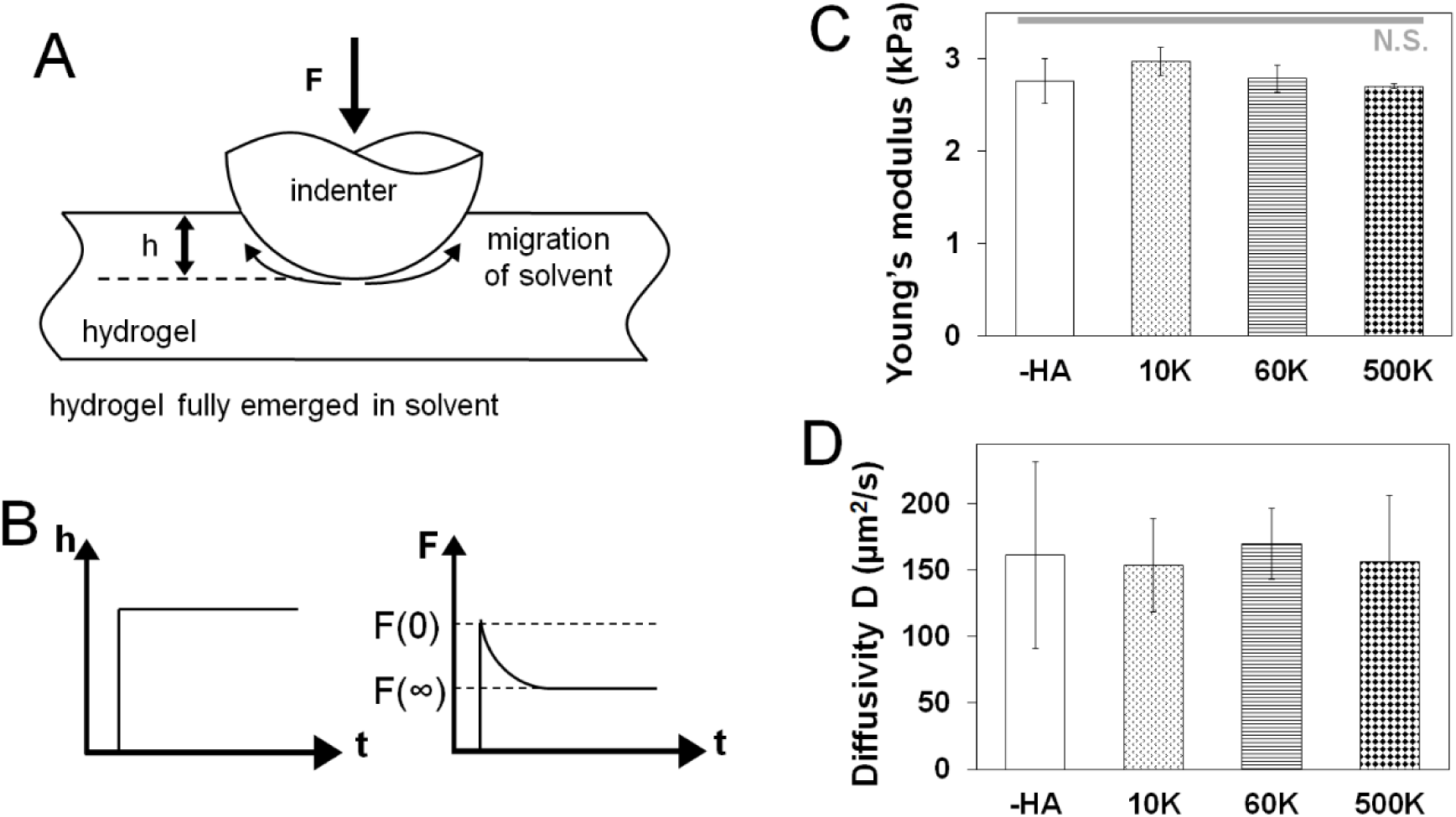
(A) Schematic drawing of measuring hydrogel water diffusivity via AFM. (B) Poroelastic parameters are extracted via indentation performed to a fixed depth followed by force relaxation to a new equilibrium state. Characterization that for a homologous series of GelMA hydrogels developed for this project there was a negligible effect of the molecular weight of hyaluronic acid incorporated into the GelMA hydrogel on (C) hydrogel Young’s modulus measured via MTS (n=6) and (D) hydrogel diffusivity via AFM (n=6).

### 2.3. Patient derived xenograft cell culture

Short-term explant cultures derived from the GBM39 PDX model were obtained from Mayo Clinic (Rochester, Minnesota). PDX samples were mechanically disaggregated, plated on low-growth factor Matrigel coated tissue culture flasks in in standard culture media made with Dulbecco’s modified eagle medium (DMEM; Gibco) supplemented with 10% fetal bovine serum (FBS; Atlanta biologicals) and 1% penicillin/streptomycin (P/S; Lonza) at 37 °C in a 5% CO_2_ environment. Flasks were shipped by overnight expression and then used upon arrival after trypsinzation. For analysis of cell metabolic health and protein expression, GBM39 cells were homogeneously mixed with the GelMA ± HAMA solution at a density of 4×10^6^ cells/mL. Cell-seeded hydrogels were incubated in cell culture medium at 37°C, 5% CO_2_ in low adhesion well plates containing standard culture media (DMEM with 10% FBS and 1% P/S). Culture media was changed at day 3 and day 5 for all cell-containing hydrogels.

### 2.4. Time-lapse cell invasion assay using spheroids

To measure relative cell motion in the fully three-dimensional hydrogel environment, we embedded GBM spheroids into our hydrogel. A methylcellulose (MC, 12 wt% in 0.5x PBS, Sigma-Aldrich) solution was made with constant stirring at 4°C overnight, then autoclaved and kept at 4°C for storage. MC solution was then added into 96-well plate and kept at 37°C overnight to form a non-adherent MC-hydrogel layer. 10^5^ GBM cells were added to each well, placed at 37°C - 5% CO_2_ environment with constant horizontal-shaking (60 rpm) overnight to aid spheroid formation (Lee et al., 2011). Spheroids were then mixed with pre-polymer GelMA ± HAMA solution, photopolymerized and cultured following the same method previously described. Cell invasion into the hydrogel was traced throughout seven-day culture by taking images on days 0 (immediately after embedding), 1, 2, 3, 5 and 7 using a Leica DMI 400B florescence microscope under bright field. Analysis of cell invasion distance (d_i_ = r_i_-r_0_) was quantified via ImageJ using the relative radius (cell spreading shape ∼ πr_i_^2^) compared to day 0 (r_0_) using a method previously described by our group (Chen et al., 2017a).

### 2.5. Analysis of cell metabolic activity

The total metabolic activity of cell-containing hydrogels was measured immediately after hydrogel encapsulation (day 0) and then subsequently at days 3 and 7 of hydrogel culture. Metabolic activity was analyzed using a dimethylthiazol-diphenyltetrazolium bromide assay (MTT; Molecular Probes) following manufacturer’s instructions. Briefly, at each time point the culture media surrounding each hydrogel sample was replaced with MTT-containing media and incubated for 4 hours, then solution was replaced with dimethyl sulfoxide (DMSO; Sigma-Aldrich) and set overnight. Metabolic activity of samples was measured via absorbance at 540 nm using a microplate reader (Synergy HT, Biotek), with data normalized to day 0 samples (immediately after seeding) as fold change.

### 2.6. Quantification of soluble hyaluronic acid secretion

The concentration of soluble HA in the media was quantified from sample media using an enzyme-linked immunosorbent assay (ELISA, R&D systems) following the manufacturer’s instructions. Sample media were collected at days 3, 5 and 7. Samples were analyzed via a microplate reader (Synergy HT, Biotek) with 450/540 nm wavelength absorbance. Soluble HA concentration within the media at each time point was calculated, with accumulated results reported as a function of all previous time point measurements.

### 2.7. Protein isolation and Western blotting

Procedures of protein isolation and Western blotting were described in previous publication (Caliari et al., 2015). Protein isolation was done by extracting proteins from cell-containing hydrogels by using cold RIPA buffer and incubating for 30 minutes. Total protein concentration in the lysates was determined by Pierce™ BCA Protein Assay Kit (Thermo Scientific). Lysates were then mixed with 2x Laemmli Sample Buffer (Bio-Rad) and 2-Mercaptoethanol (Sigma-Aldrich), heated to 95°C for 10 minutes, then loaded (3 μg protein loaded onto per lane) onto polyacrylamide gels (4%-20% gradient; Bio-Rad). Gel electrophoresis was performed at 150 V. Proteins were then transferred onto nitrocellulose membrane (GE Healthcare) using Trans-Blot SD (Bio-Rad) under 300 mA for 2 h. Membranes were then cut into desired MW range and blocked in blocking buffer for 1 h followed by primary antibodies incubation at 4 °C overnight. Membranes were subsequently washed with Tris Buffered Saline with Tween20 (TBST), followed by secondary antibody incubation for 2 hours at room temperature. Imaging signal was visualized using imaging kits (SuperSignal™ West Pico PLUS Chemiluminescent Substrate or SuperSignal™ West Femto Maximum Sensitivity Substrate, Sigma-Aldrich) via an Image Quant LAS 4010 chemiluminescence imager (GE Healthcare). Band intensities were quantified using ImageJ and normalized to β-actin expression. Buffers and antibodies used in each condition are listed (**Table S1**).

### 2.8. Statistics

All statistical analysis was performed using one-way analysis of variance (ANOVA) followed by Tukey’s test. A minimum sample number of n = 3 (MTT, ELISA, Western), n = 6 (Young’s modulus, diffusivity, invasion) samples were used for all assays. Statistical significance was set at p < 0.05. Error is reported as the standard error of the mean.

## 3. Results

GelMA hydrogels lacking matrix-bound HA will be denoted as “-HA” while hydrogels containing 15 w/w% HAMA will be denoted as “10K”, “60K”, or “500K” to denote the molecular weight of the incorporated HA sodium.

### 3.1. Molecular weight of matrix-bound HA does not impact Young’s moduli or diffusive properties of the family of gelatin hydrogels

The biophysical properties of the homologous series of GelMA hydrogel (-HA, 10K, 60K, 500K) were assessed via unconfined compression and AFM indentation. The Young’s moduli of all hydrogels did not vary as a result of inclusion of matrix-bound HA regardless of the HA MW. Critically, the Young’s modulus of these hydrogels (-HA: 2.76 ± 0.24 kPa; 10K: 2.97 ± 0.15 kPa; 60K: 2.79 ± 0.15 kPa; 500K: 2.70 ± 0.03 kPa) are within physio-relevant range (10^0^-10^1^ kPa) for the GBM TME. Similarly, the diffusivity of all hydrogel variants was not significantly influenced by the presence or absence of matrix immobilized HA (-HA: 161.04 ± 70.33 µm^2^/s; 10K: 153.54 ± 34.92 µm^2^/s; 60K: 169.90 ± 26.88; 500K: 156.43 ± 50.18 µm^2^/s) (Figure 1).

### 3.2. Metabolic activity of GBM39 PDX cells cultured in GelMA hydrogels is sensitive to the molecular weight of matrix bound HA

The metabolic activity of GBM39 PDX cells encapsulated within the homologous series of GelMA hydrogels (-HA, 10K, 60K, 500K) was traced through 7 days in culture, with results normalized to day 0 values for each group. The groups with matrix-bound HA (10K, 60K and 500K) showed a significantly higher metabolic activity compare to -HA group (p < 0.05), with the 60K HA group showing the highest metabolic activity amongst all groups (Figure 2).

**Figure 2.**
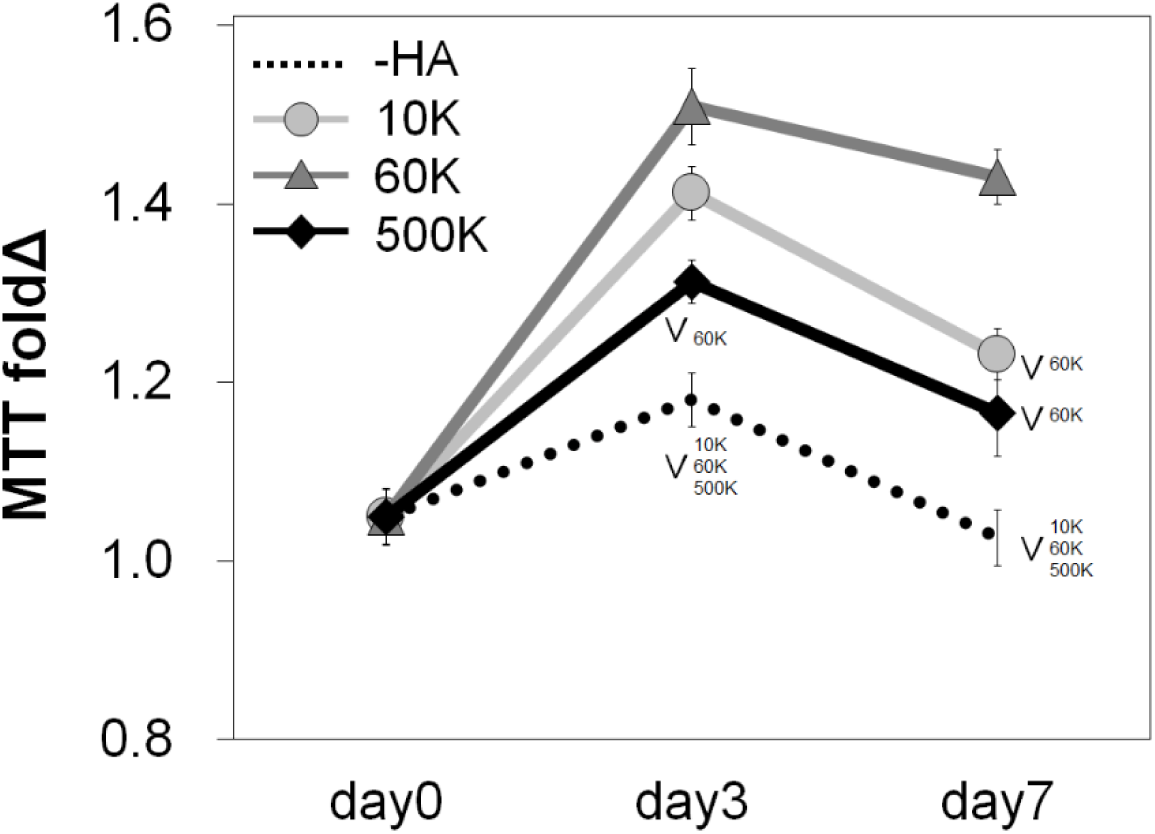
Overall metabolic activities (n=3) of GBM seeded GelMA hydrogels as a function of incorporated hyaluronic acid molecular weight. Results are provided throughout the 7-day culture and are normalized to the metabolic activity of each group at day 0. Samples containing matrix-bound HA showed an overall higher metabolic activity compare to GelMA only (–HA) hydrogels. The greatest metabolic activity was observed for GelMA hydrogels containing 60 kDa (60K) HA. ^∨^ p < 0.05 significant decrease between groups.

### 3.3. The molecular weight of matrix-bound HA significantly affects invasion

The invasion of GBM39 PDX cells into the surrounding hydrogel matrix was measured via a previously reported spheroid assay through 7-days in culture. GBM39 invasion was strongly influenced by hydrogel HA content. The highest level of invasion was observed for GelMA hydrogels either lacking matrix bound HA (-HA), or those containing mid-range (60K) molecular weight matrix-immobilized HA (Figure 3). At early-to-mid time points (up to day 5), GBM cell invasion was significantly depressed in the low molecular weight 10K group, but GBM invasion increased steeply at later time points (day 7), matching the highest invasion groups. GelMA hydrogels containing the largest molecular weight HA (500K) showed significantly reduced invasion compared to all other hydrogel groups (-HA, 10K, 60K) throughout the entire period studied.

**Figure 3.**
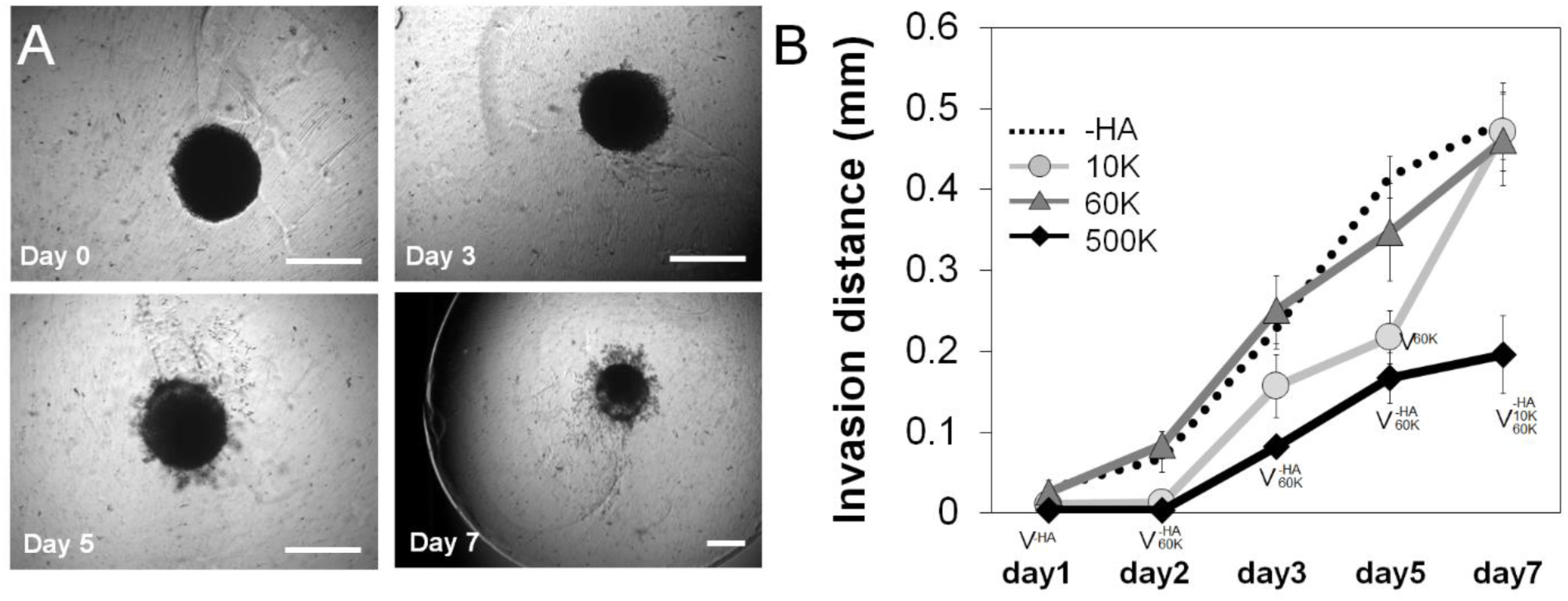
(A) PDX invasion (n=6) into the surrounding hydrogel was quantified via a spheroid assay throughout the 7-day culture period. Representative images of spheroid invasion throughout the seven day culture, showing GBM cells progressively leave the spheroid and invade the hydrogel. Scale bar 0.5 mm. (B) Quantification of GBM cell invasion into the GelMA hydrogel as a function of the molecular weight of matrix immobilized HA. GelMA only (-HA) and GelMA hydrogels containing 60 kDa (60K) HA showed the greatest levels of invasion, with no significant difference between those groups across the culture period. Interestingly, GelMA hydrogels containing high molecular weight HA (500K) showed significantly reduced invasion. ^∨^ p < 0.05 significant decrease between groups.

### 3.4. The accumulation of soluble HA in media reflects matrix-composition

ELISA was performed to measure the concentration of soluble HA in the culture media over the course of the invasion experiment. An increase in soluble HA concentration was observed in the hydrogels lacking matrix bound HA (-HA) compared to all groups containing matrix-bound HA. Interestingly, the presence of soluble HA for hydrogel groups containing matrix-immobilized HA was found to be strongly associated with the molecular weight of immobilized HA, with 500K group showing significantly upregulated secretion compared to GBM cells in 10K and 60K HA hydrogels as early as day 3. Significant increases were observed in soluble HA production in 60K vs. 10K hydrogels appeared by day 7 of culture (Figure 4).

**Figure 4.**
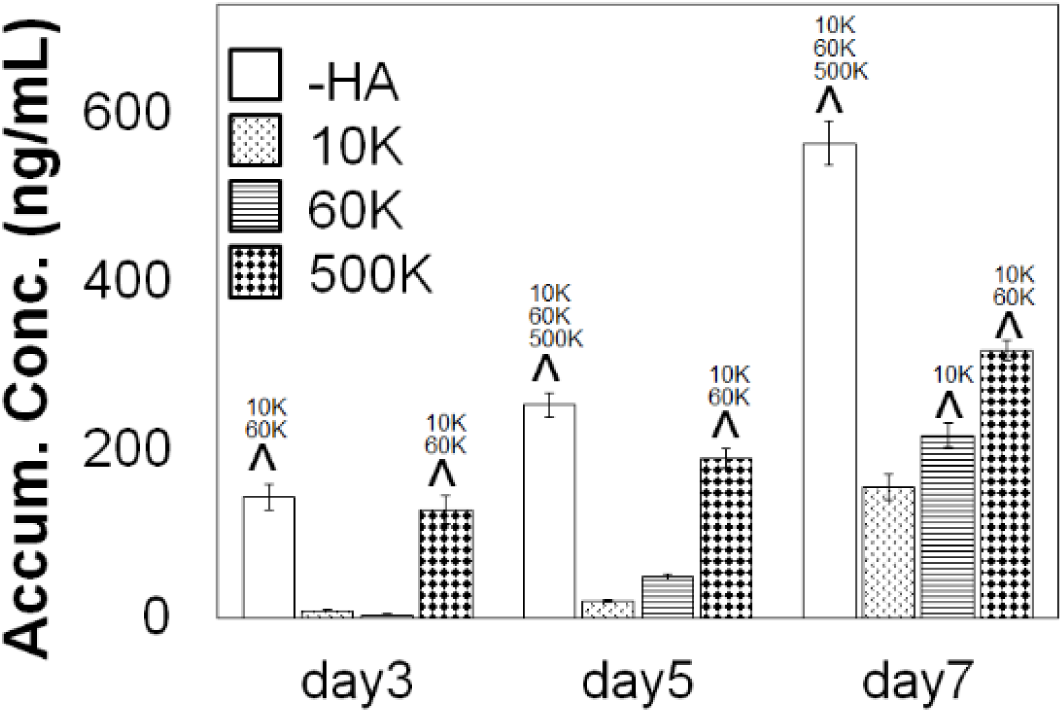
Accumulative of soluble HA in the media over the course of GBM culture in GelMA hydrogels, measured via ELISA (n=3). GBM cells in GelMA hydrogels lacking any matrix immobilized HA (-HA) showed secreted significantly higher amount of soluble HA compare to GBM cells cultured within GelMA hydrogels containing matrix-bound HA. Production of soluble HA by GBM cells in GelMA hydrogels containing matrix-immobilized HA were strongly sensitive to the molecular weight of the matrix immobilized HA. Notably, soluble HA secretion increased with the MW of immobilized HA, with the 500K group secreting significantly higher amount of soluble HA compare to 60K and 10K. ^ p < 0.05 significant increase between different groups.

### 3.5. Protein expression of hyaluronic acid remodeling associated proteins were not strongly influenced by hydrogel HA content

The expression of protein families, biosynthetic hyaluronan synthase (HAS1, HAS2, HAS3) and degradative hyaluronidase (HYAL1, HYAL2), associated with HA remodeling were subsequently quantified via Western blot analyses (Figure 5, **Figure S1-S2**). No significant differences were observed in expression levels within each group as a function of immobilized HA molecular weight. However, GBM cells in the highest molecular weight HA hydrogels (500K) showed generalized increases in both HAS and HYAL (significant for HYAL2) compared to all other hydrogel conditions.

**Figure 5.**
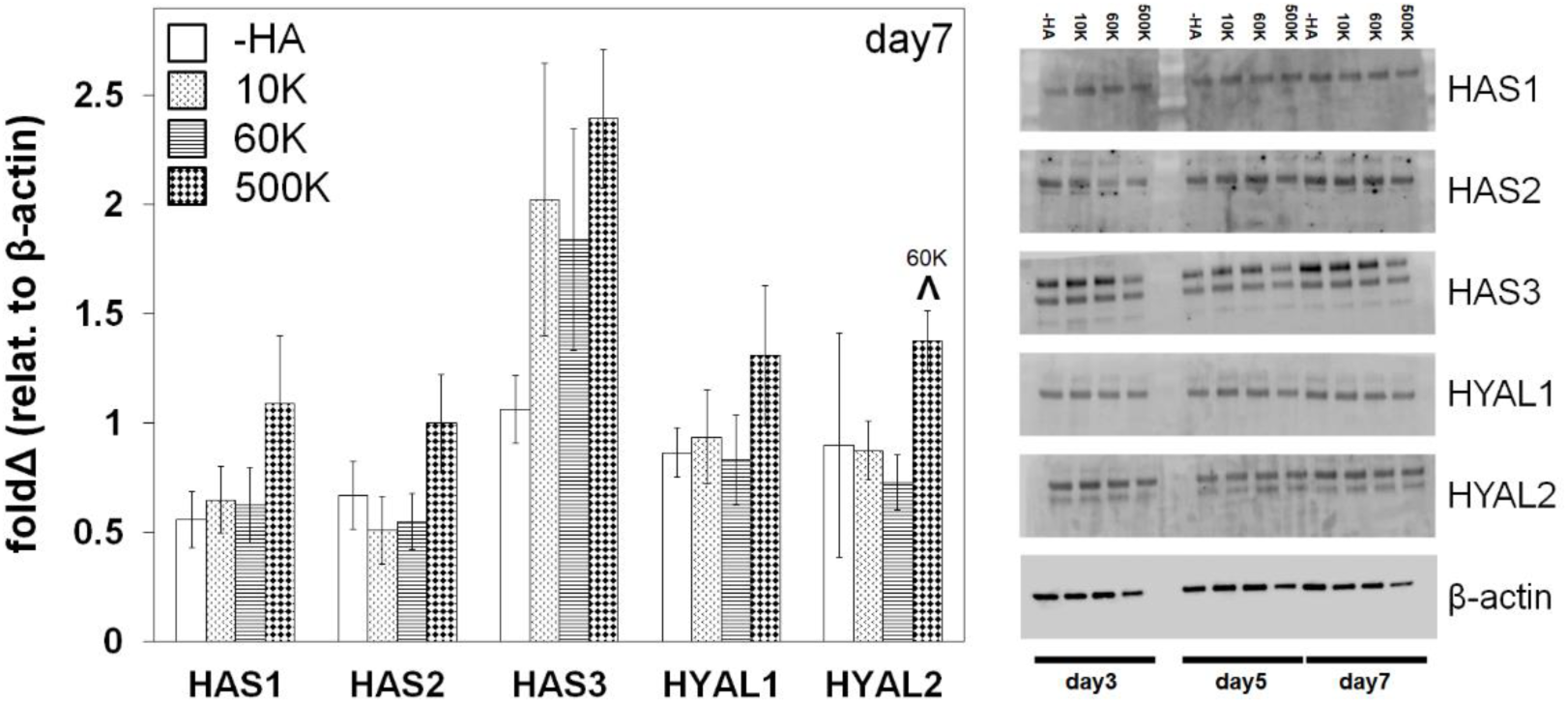
Hyaluronan synthase (HAS) and hyaluronidase (HYAL) protein expression of GBM cells in gelatin hydrogels as a function of matrix immobilized HA molecular weight, analyzed via Western Blot at day 7 (n=3). β-actin is used as loading control. ^ p < 0.05 significant increase between different groups.

## 4. Discussion

The heterogeneity of GBM tumor microenvironment complicates its study both in vivo and in vitro. Within that high diversity, the extracellular HA has been widely associated with cancer invasion and response to treatment (Park et al., 2008;Rankin and Frankel, 2016;Zhao et al., 2017). Naturally, HA is synthesized and deposited in the extracellular space by HAS family and degraded into different size fragments by HYAL enzyme family. The alteration of the levels of these enzymes are associated with various types of diseases. LMW HA (< 30 kDa) has been associated mainly with increased tumor growth, cell migration and angiogenesis, while HMW (250 to >1000 kDa) is commonly believed to lead to greater structural stability with reduced tumor growth, migration, and angiogenesis (Monslow et al., 2015). However, despite their relevance in GBM microenvironment, the influence of HA MW has been largely neglected in regard to the construction of ex vivo biomaterial platforms to examine GBM cell activity. This project seeks to understand the effect of HA molecular weight, both matrix bound and cell secreted, on the invasive phenotype of a patient-derive GBM specimen. We developed and characterized a homologous series of HA-decorated gelatin-based hydrogels to evaluate the effect of HA MW on GBM invasiveness and phenotypic responses.

A family of hydrogels with no matrix-bound HA or with increasing MW HA (10kDa, 60kDa and 500kDa) was fabricated using a method previously described (Pedron et al., 2013b;Chen et al., 2017c). Studies demonstrate that substrate stiffness and diffusion can deeply influence the migration capacity of GBM cells in HA containing hydrogels (Rape et al., 2014;Umesh et al., 2014;Wang et al., 2014;Chen et al., 2017b). However, we have previously described a framework to adjust the relative ratio of GelMA to HA content as well as manipulating the crosslinking conditions to generate a series of GelMA hydrogels containing increasing wt% of a single MW HA (Pedron et al., 2013a). We therefore adapted this approach to create the homologous series of hydrogels described in this study, that contained a conserved wt% of HA but that varied the MW of matrix-immobilized HA. We then employed a series of biophysical and biochemical characterization protocols to describe poroelastic features of these hydrogels. Crosslinking density can be preserved by adjusting the photoinitiator concentration in the pre-polymer solution (Table 1), and therefore maintaining the Young’s modulus between different hydrogels. Moreover, the deformation of the gel in contact with the AFM tip results from two simultaneous molecular processes: the conformational change of the network, and the migration of the solvent molecules (Hu et al., 2010). In this case, the poroelasticity of the hydrogels, characterized by the diffusivity (Figure 1D), stays unchanged for all samples used. Both Young’s modulus and diffusivity showed no significant difference among all groups suggesting these hydrogels were able to provide similar culture conditions for cells while providing the opportunity to adjust the molecular weight of bound HA.

We subsequently measured the metabolic activity of GBM39 PDX cells as a function of matrix bound HA MW. The presence of matrix-bound HA aided GBM metabolic response compared to the -HA group. In general, all cells remained viable within the hydrogel up to 7 days, without showing apoptosis or cell death. Further, we performed a spheroid-based invasion assay to investigate the effects of matrix-bound HA MW on invasion at different time points, including early (1 and 2 days), mid (3 and 5 days) and longer (7 day) time points. Consistent with earlier observations described by our group using GBM cell lines (Chen et al., 2017a;Chen et al., 2018), we found GBM invasion in GelMA hydrogels lacking matrix bound HA was greatest. However, invasion was strongly influence by the MW of immobilized HA with GBM cell invasion in hydrogels containing 60kDa being equivalent to hydrogels lacking matrix bound HA. Further, this invasive potential of GBM39 cells within –HA and +HA hydrogels is not associated to their metabolic activity profiles (Figure 2). Although migration and proliferation are considered to be circumscribed phenotypes that do not co-occur with each other in GBM, the complex microenvironment of PDX suggests that both can coexist. Moreover, GBM cells adapt to the different phenotypes by using regulatory signaling from the local microenvironment (Xie et al., 2014). Interestingly, while invasion was initially significantly reduced in low MW HA hydrogels (10K), GBM invasion increased significantly at later time points. However, GBM invasions was strongly reduced in GelMA hydrogels containing high molecular weight HA (500K) throughout the entirety of the study, suggesting more mature HA matrices will inhibit GBM invasion. While recent studies have begun to examine the design of implants to reduce GBM invasion (Jain et al., 2014), these findings suggest an interesting line of future studies that wound center on the incorporation of hydrogels into the resection margins containing attractive biomechanical and biomolecular properties to potentially recruit nearby GBM cells as a means to reduce invasive spreading. Regardless, the presence of both fibrillar and HA associated features of the TME in these HA decorated GelMA hydrogels may be particularly useful in the context of GBM invasion in perivascular niches that contain such matrix diversity (Ngo and Harley, 2018).

Studies have shown that HMW HA could inhibit tumor invasion by inhibiting MMPs production and down-regulating invasion related pathways such as MAPK and Akt (Chang et al., 2012), while LMW HA may promote these invasion related pathways (West et al., 1985;Lam et al., 2014). We hypothesized that the significant decrease of motility in PDX cells in 500kDa hydrogels is due to the down-regulation of invasion related pathways, induced by the local extracellular microenvironment. We observed endogenous HA production was significantly elevated without the presence of matrix-bound HA (-HA) (Figure 4), consistent with previous studies reported by our group using immortalized cell lines that demonstrated soluble HA production was associated with increased GBM cell invasion (Chen et al., 2017c). More interestingly, soluble HA production across the homologous series of hydrogels tested in this study (-HA, 10K, 60K, 500K) showed greatest endogenous HA production in hydrogels lacking matrix immobilized HA. However, endogenous production of HA was also sensitive to the molecular weight of matrix bound HA, with greater endogenous HA production seen with increasing molecular weight of bound HA. This trend of increasing soluble HA production with increasing molecular weight of matrix-bound HA may be associated to an adaptation required to mobilize matrix bound HA for invasion.

Many studies have shown that the levels of HAS correlate with breast and colon cancer malignancy and patient prognosis (Bullard et al., 2003;Auvinen et al., 2014). Inhibition of HAS has been used as an alternative therapeutic strategy using mRNA silencing HAS or HAS-targeting drugs (e.g. 4-Methylubelliferone) (Nakamura et al., 1997;Li et al., 2007;Nagy et al., 2015). While some studies suggest addition of HYAL into chemotherapy efficiently improves the patient prognosis (Baumgartner et al., 1998;Klocker et al., 1998;Stern, 2008), others show HYAL levels are correlated with cancer malignancy and invasiveness in breast, prostate and bladder cancers (Lokeshwar et al., 1996;Madan et al., 1999;Lokeshwar et al., 2005;Stern, 2008). While we observe no significant across-the-board trends in HAS and HYAL proteins levels as a function of matrix immobilized HA, GBM cells in hydrogels containing the highest molecular weight HA (500 kDa) show overall a higher expression of all HAS and HYAL families compared to the rest. However, these results did not directly correlate with the GBM invasiveness as for what we observed. While HAS and HYAL both play a key role in tumor progression and invasiveness, the dynamic balance might be more crucial instead of one over the other, suggesting opportunities for future studies using an expanded library of patient-derived GBM specimens using this homologous series of GelMA hydrogels.

High production of HA is normally associated with tumor progression, although overly high levels of hyaluronic acid secretion may lead to an opposite behavior (Itano et al., 2004). Moreover, in gliomas, this HA associated tumor progression only occurs if hyaluronan is expressed simultaneously with HAS (Enegd et al., 2002). Therefore, studies suggest that HA turnover is required for the increase of HA associated GBM tumor malignancy. Additionally, the relative contribution of matrix-bonded and cell produced HA increases this complexity. Therefore, a feedback mechanism between stromal and produced HA has been drafted for epithelial cancers (Koyama et al., 2007) but is still unexplored in glioblastoma. In this study using an ex vivo biomaterial model, we show how the dynamic interplay between extracellular matrix associated and cell produced HA affects GBM cell behavior. Further ongoing research may allow identification of alternative antitumor treatments in the context of the GBM microenvironment.

## 5. Acknowledgements

Hydrogel diffusivity measurements were carried out in part in the Frederick Seitz Materials Research Laboratory Central Research Facilities, University of Illinois at Urbana-Champaign. HA MW distribution imaging was carried out in the Protein Sciences center of the Roy J. Carver Biotechnology Center, University of Illinois at Urbana-Champaign. Research reported in this publication was supported by the National Cancer Institute, National Institute of Diabetes and Digestive and Kidney Diseases, and the National Institute of Biomedical Imaging and Bioengineering of the National Institutes of Health under Award Numbers R01CA197488 (BACH), R01 DK099528 (BACH), and T32EB019944 (JEC). The content is solely the responsibility of the authors and does not necessarily represent the official views of the National Institutes of Health. The authors are also grateful for additional funding provided by the Department of Chemical & Biomolecular Engineering (BACH) and the Carl R. Woese Institute for Genomic Biology (BACH) at the University of Illinois at Urbana-Champaign. Development and maintenance of the GBM PDX models was supported by Mayo Clinic, the Mayo SPORE in Brain Cancer (CA108961), and the Mayo Clinic Brain Tumor Patient-Derived Xenograft National Resource (NS092940).

## 7. Author Contributions

JEC, SP and BACH designed experiments, performed cell experiments, data analysis, results interpretation, and wrote the manuscript. PS, YH performed AFM experiments and assisted with manuscript writing. JS assisted with experiment design, results interpretation, and manuscript writing. BACH is the principal investigator.

